# RCoV19: A One-stop Hub for SARS-CoV-2 Genome Data Integration, Variants Monitoring, and Risk Pre-warning

**DOI:** 10.1101/2023.09.24.558358

**Authors:** Cuiping Li, Lina Ma, Dong Zou, Rongqin Zhang, Xue Bai, Lun Li, Gangao Wu, Tianhao Huang, Wei Zhao, Enhui Jin, Yiming Bao, Shuhui Song

**Affiliations:** National Genomics Data Center, Beijing Institute of Genomics, Chinese Academy of Sciences and China National Center for Bioinformation, Beijing 100101, China; CAS Key Laboratory of Genome Sciences and Information, Beijing Institute of Genomics, Chinese Academy of Sciences and China National Center for Bioinformation, Beijing 100101, China; University of Chinese Academy of Sciences, Beijing 100049, China; Sino-Danish College, University of Chinese Academy of Sciences, Beijing 100049, China

**Keywords:** SARS-CoV-2, Mutation, Variants, Surveillance, Pre-warning

## Abstract

The Resource for Coronavirus 2019 (RCoV19, https://ngdc.cncb.ac.cn/ncov/) is an open-access information resource dedicated to providing valuable data on the genomes, mutations, and variants of the severe acute respiratory syndrome coronavirus 2 (SARS-CoV-2). In this updated implementation of RCoV19, we have made significant improvements and advancements over the previous version. Firstly, we have implemented a highly refined genome data curation model. This model now features an automated integration pipeline and optimized curation rules, enabling efficient daily updates of data in RCoV19. Secondly, we have developed a global and regional lineage evolution monitoring platform, alongside an outbreak risk pre-warning system. These additions provide a comprehensive understanding of SARS-CoV-2 evolution and transmission patterns, enabling better preparedness and response strategies. Thirdly, we have developed a powerful interactive mutation spectrum comparison module. This module allows users to compare and analyze mutation patterns, assisting in the detection of potential new lineages. Furthermore, we have incorporated a comprehensive knowledgebase on mutation effects. This knowledgebase serves as a valuable resource for retrieving information on the functional implications of specific mutations. In summary, RCoV19 serves as a vital scientific resource, providing access to valuable data, relevant information, and technical support in the global fight against COVID-19.

## Introduction

SARS-CoV-2 is responsible for the COVID-19 pandemic, and continues to evolve and spread to threat public health worldwide. Genome data play a crucial role in understanding mutations (refers to an actual nucleotide or amino acid change in a viral genome), functions, and supporting the design of candidate vaccines. While there are various data deposition repositories available, such as EpiCoV^TM^ [1], GenBank [2, 3], and GenBase (https://ngdc.cncb.ac.cn/genbase/), none of them encompass all worldwide genome data, and redundancies exist among these repositories. Therefore, the need for a comprehensive SARS-CoV-2 database arises to integrate genome data, monitor evolution, and provide pre-warnings for high-risk variants. Such a database is essential to comprehend the ongoing pandemic and facilitate timely adjustments to public health interventions.

With millions of genome sequences now available, several platforms have been developed to track SARS-CoV-2 mutations. These platforms, including COVID-19 CG [4], Outbreak [5], and CoV-Spectrum [6], enable tracking of mutations by location, date of interest, and known variants globally. VarEPS [7] assesses the risk level of mutations and variants based on their transmissibility and affinity to neutralizing antibodies. Additionally, databases like CoV-RDB [8, 9] and COG-UK-ME [10] have compiled mutations associated with reduced susceptibility to various factors, such as clinical stage SARS-CoV-2 Spike monoclonal antibody (mAb), RNA-dependent RNA polymerase (RdRP) inhibitor, 3C-like protease (3CLpro) inhibitor, or mutations on T cell epitope. However, despite these significant efforts, there are limitations in terms of efficiency and comprehensiveness. Most of these platforms and databases only focus on specific aspects of SARS-CoV-2 monitoring or prevention.

Furthermore, numerous important mutations affecting transmissibility, infectivity, or expression are scattered throughout published literature. Consequently, there is an urgent need to build an integrated and comprehensive system that encompasses “data-information-knowledge-application”. This system should provide real-time services for sequence monitoring, evolution tracking, and pre-warning of high-risk variants.

RCoV19, previously known as 2019-nCoVR [11, 12], is an open-access information resource for SARS-CoV-2. It has been available online and has already provided data services to over 3.2 million visitors from 182 countries/regions worldwide, with more than 14 billion data downloads in total. In this updated release of RCoV19, significant improvements have been made in data curation, integration, sequence growth and lineage evolution surveillance, and mutation comparisons of sequences and lineages. Additionally, a weekly report on potentially high-risk haplotypes (a distinct virus genome sequence) and variants (a viral genome that may contain one or more mutations, which may affect virus’s properties) is provided, combining genetic mutation effects and haplotype network features [13, 14]. Furthermore, RCoV19 curates an integrated knowledge of mutation effects from literatures and databases, offering critical insights into virus evolution, immune escape, and medical countermeasures. Ultimately, RCoV19 establishes a one-stop hub for SARS-CoV-2 genome data integration and variant monitoring, as illustrated in **Figure 1**.

**Figure 1.**
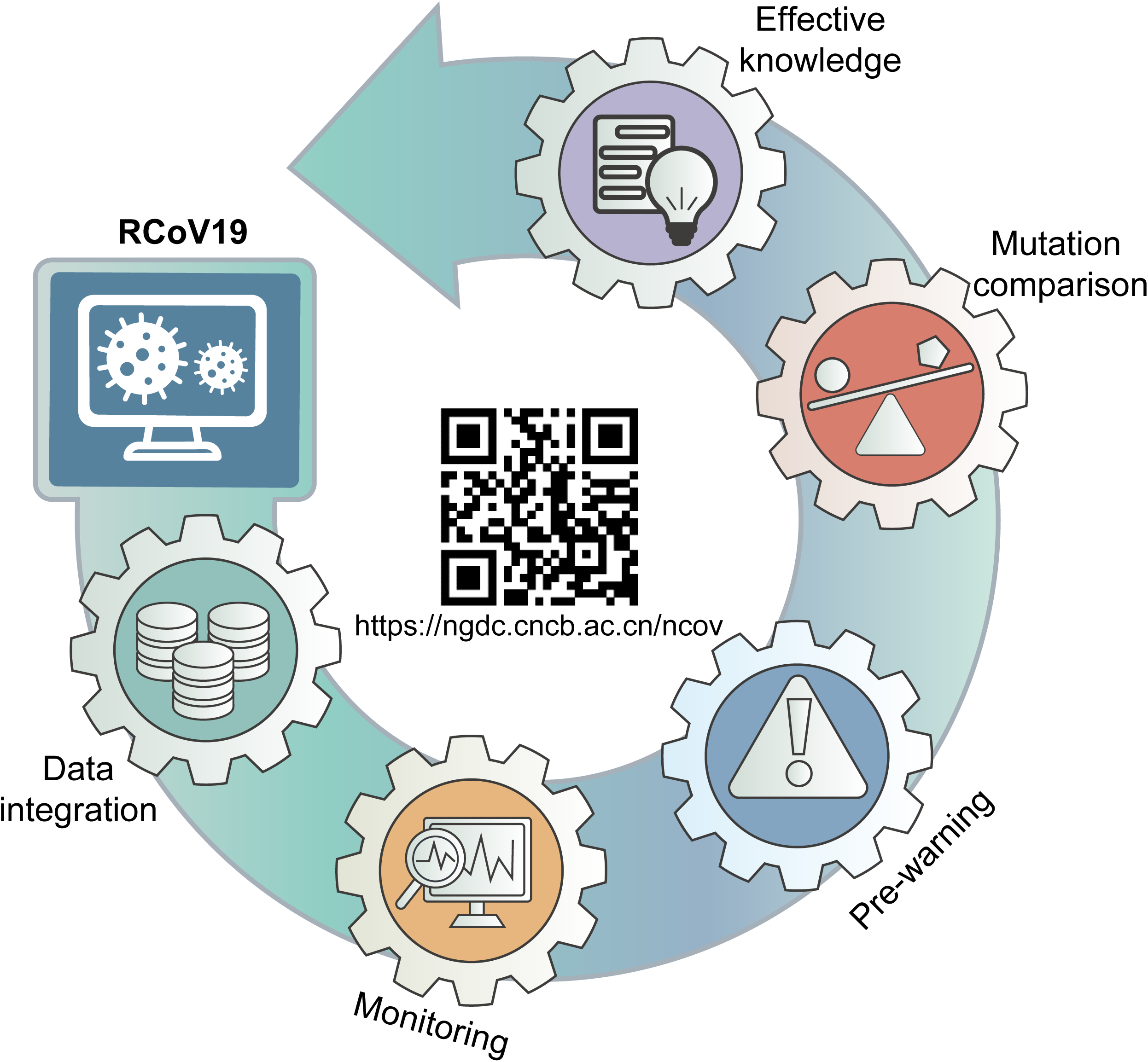
Logical architecture diagram of RCoV19 database.

## Database content and features

### Efficient integration and retrieval of worldwide SARS-CoV-2 genome data

RCoV19 is an extensive data resource for SARS-CoV-2 that collects genome data from multiple repositories, performs de-redundancy processing, and assesses sequence quality to ensure a comprehensive and curated collection of worldwide genomes (**Figure 2**). The resource incorporates data from repositories such as EpiCoV^TM^ [1], GenBank [2, 3], CNGBdb [15], and Novel Coronavirus Service System of NMDC [16], and has included data from GenBase since the beginning of 2023. To eliminate redundancies, RCoV19 identifies identical genomes across different sources and cross-references related accession IDs. Notably, 91.3% GenBank sequences overlap with EpiCoVTM sequences, while 56.7% of EpiCoVTM sequences are unique (**Figure S1**). It determines completeness of the protein-coding region, evaluates sequences in five aspects (Ns, degenerate bases, gaps, mutations, and mutation density) and defines high-quality sequences based on Ns and degenerate bases. These processes enable RCoV19 to provide a comprehensive and reliable list of SARS-CoV-2 genomes for global monitoring and pre-warning purposes.

**Figure 2.**
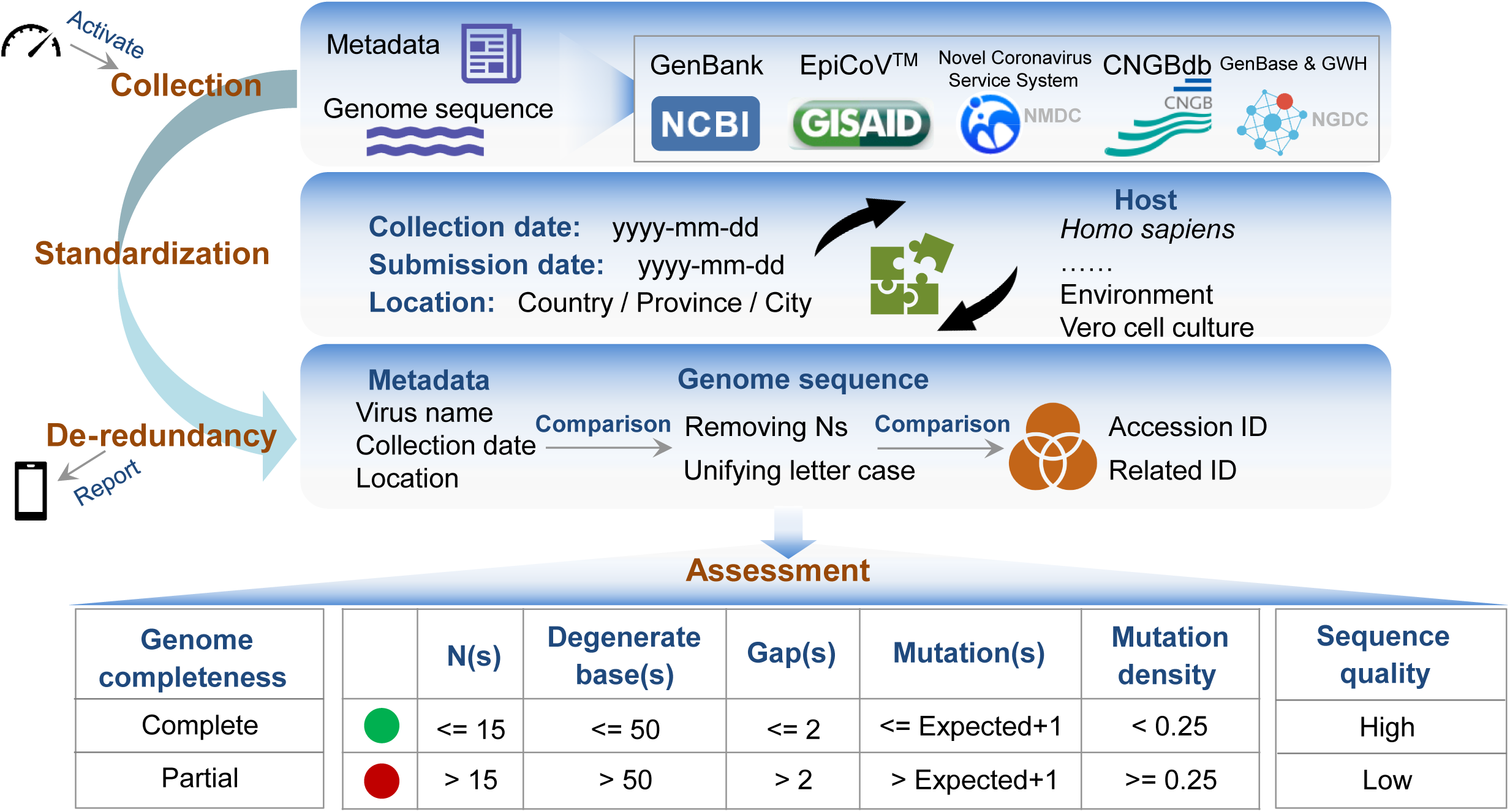
Framework of genome data curation model for SARS-CoV-2. RCoV19 integrates genome data from different repositories and provides valued-added curations. It collects metadata and genome sequences from different resources, standardizes metadata, and performs de-redundancy processing based on metadata and sequence comparison. These steps have been chained together as one workflow, which is activated automatically every day and sends the integration statistics to mobile phone client at the end. After integration, RCoV19 performs a series of assessments; it determines completeness of the protein-coding region, assesses sequence quality in five aspects, and defines high-quality sequences. We consider a sequence to be of high quality if it could pass quality control for both Ns (<= 15) and degenerate bases (<= 50). Otherwise, it is of low quality.

In the new version, the SARS-CoV-2 genome data curation model has been significantly enhanced with an automated integration pipeline and optimized curation rules (**Figure 2**), ensuring efficient daily updates in RCoV19. The automated pipeline, activated by a timer every day, collects genome data from various repositories through the Chrome Browser on Linux, standardizes genome metadata, and performs de-redundancy processing. This automated approach improves efficiency compared to semi-automated methods and enables regular and constant updates. Curation rules have also been optimized to achieve more accurate de-redundancy, by comparing genome sequences (with removal of Ns and uniform letter case) in addition to key metadata (virus name, sampling date and location). Furthermore, the curation rule for assessing abnormally high mutations has been improved. The expected number of mutations for each sequence is now calculated based on its sampling date and empirical mutation rate [17], providing a more realistic assessment. If the observed number of mutations exceeds the expected number, the genome sequence is highlighted with a red dot, indicating the need for further investigation into sequencing quality issues.

With the automated integration pipeline and optimized curation rules, RCoV19 accommodated a total of 16,119,080 non-redundant genome sequences from 193 countries/regions as of June 10, 2023. A comprehensive and up-to-date list of all released SARS-CoV-2 genome metadata can be freely accessed and downloaded by users at https://bigd.big.ac.cn/ncov/release_genome. The majority of these genomes are contributed by countries such as the United States (31.6%), United Kingdom (19.3%), Germany (5.9%), France (4.4%), Denmark (4.0%), Japan (3.8%), and Canada (3.4%). Among the released human-derived genome sequences (16,103,219), 87.7% are complete, and 47.0% are both complete and high-quality. Additionally, RCoV19 offers the service of collapsing identical sequences, resulting in a total of 5,832,804 unique sequences (1:1.3) among the complete and high-quality human-derived genome sequences, and 13,762,271 unique sequences (1:1.2) among all released genomes, highlighting the rapid evolution and high diversity of SARS-CoV-2 genomes.

To facilitate fast and customized retrieval of SARS-CoV-2 genomes from this vast collection, RCoV19 has developed an advanced search module at https://ngdc.cncb.ac.cn/ncov/genome/search. Users can query by accession ID, Pango lineage, WHO variant label, country/region, host, nucleotide completeness, quality assessment, database resource, sampling date, and sequence length range. The search results are complemented by statistics displayed on the right side of the search page, showcasing distributions in nucleotide completeness, sequence quality, data source, WHO variant label, lineage, country/region, and host. Furthermore, all filtered results can be easily downloaded to support downstream analysis.

### Timely monitoring of sequence growth and lineage evolution

With the rapid accumulation of SARS-CoV-2 genome sequences, the emergence of new lineages in specific regions or the whole world has become increasingly prevalent. To enhance our understanding of SARS-CoV-2 evolution and transmission characteristics, we have developed specific modules for monitoring global and regional sequence growth and lineage evolution.

Sequence growth serves as an indicator of a country’s monitoring capability and level. By examining the cumulative curve of genome sequence growth based on release dates, we can identify three distinct periods: slow growth (January 2020 to March 2021), fast growth (April 2021 to April 2022), and relatively slow growth (May 2022 to present) (**Figure 3A**). We dynamically display the sequence numbers for the top ten countries each month to visualize their contributions (**Figure 3B**). Moreover, we organize sequence numbers for each country/region in a tabular format to provide various detailed data (**Figure 3C**). For example, as of June 10, 2023, a total of 67,149 sequences have been released for China (include Taiwan, HongKong and Maco), with an average release rate of dozens of sequences per day in May 2023.

**Figure 3.**
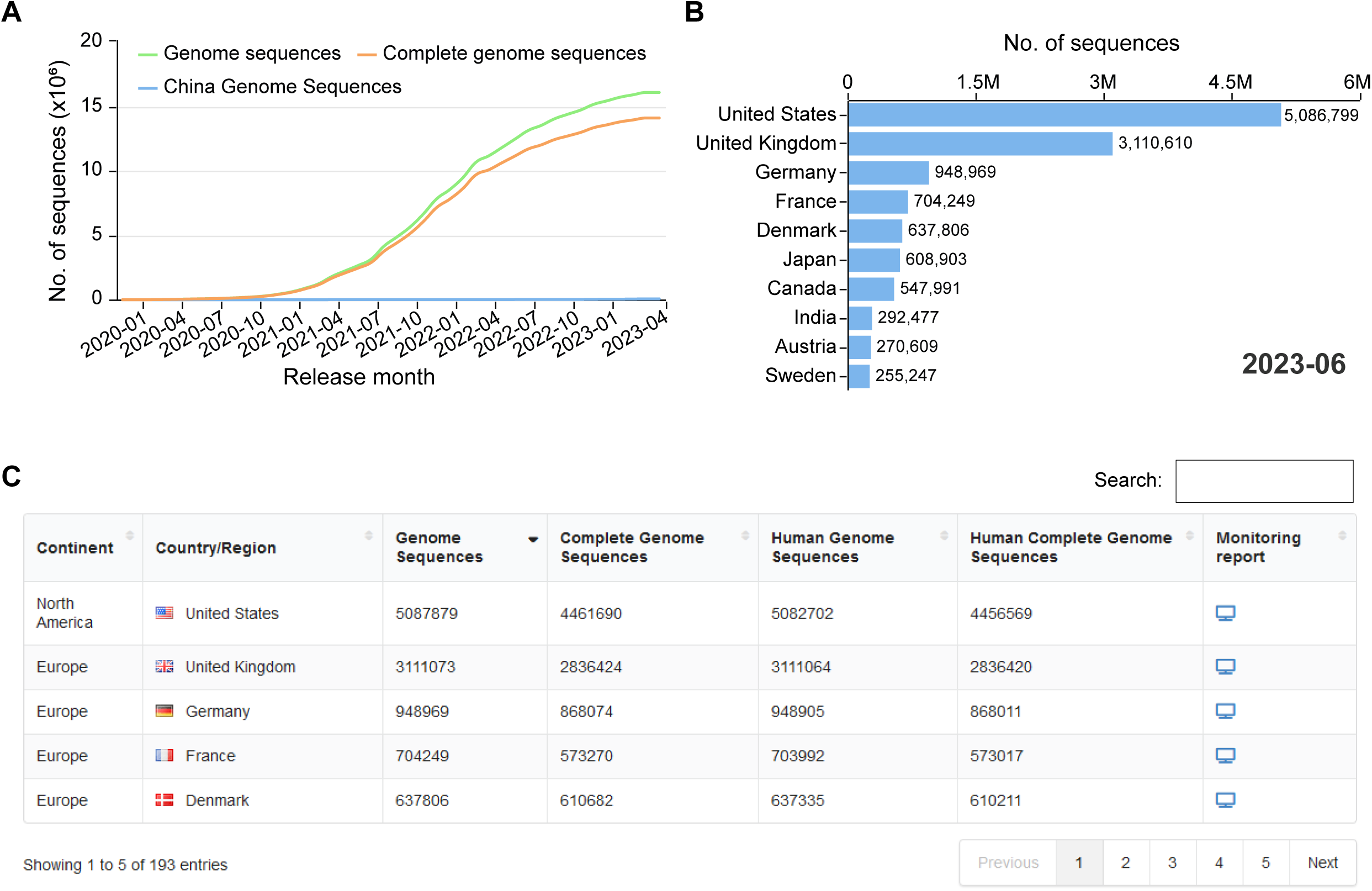
The monitoring platform of SARS-CoV-2 sequence growth globally and regionally. **A**. The dynamic growth curve of globally and China released genome sequences, and globally released complete genome sequences. **B**. A bar chart shows the top ten countries with the most public released sequences as of June 3, 2023. **C**. A tabular table shows the statistic of sequences in country/region.

As SARS-CoV-2 spreads, mutations constantly occur and accumulate, leading to the emergence of new lineages and variants. To monitor mutation rates, we calculate the mutation frequency (mutation numbers / genome length) for each genome and plot the daily median mutation frequency as a curve (**Figure 4A**). By observing the slope of curve growth, it is facilitated to timely monitor signals indicating accelerated mutation. For instance, the median mutation frequency rapidly increased to 2.1‰ in mid-December 2021 due to the rapid spread of Omicron variant and reached 3.28‰ in March 2023 due to the spread of XBB.1.5 variant. As sequences with similar mutation spectra are always classified into a Pango lineage [18] or named as a WHO-defined variant (https://www.who.int/news/item/31-05-2021-who-announces-simple-easy-to-say-labels-for-sars-cov-2-variants-of-interest-and-concern), we display the weekly sequence proportion for each lineage or variant. To highlight the main lineages or variants that are currently or previously popular, we interactively display only the top three Pango lineages or WHO-defined variants (**Figure 4B**). Additionally, the sequence proportion for each lineage is represented in a heat map (**Figure 4C**), providing informative insights into lineage trends over time. Taking China as an example, it experienced a wave of COVID-19 infections from late 2022 to early 2023. We developed an interactive map panel to dynamically display the sequence proportions of different lineages and monitor the prevalence and transmission of SARS-CoV-2 at the provincial level (**Figure 4D**).

**Figure 4.**
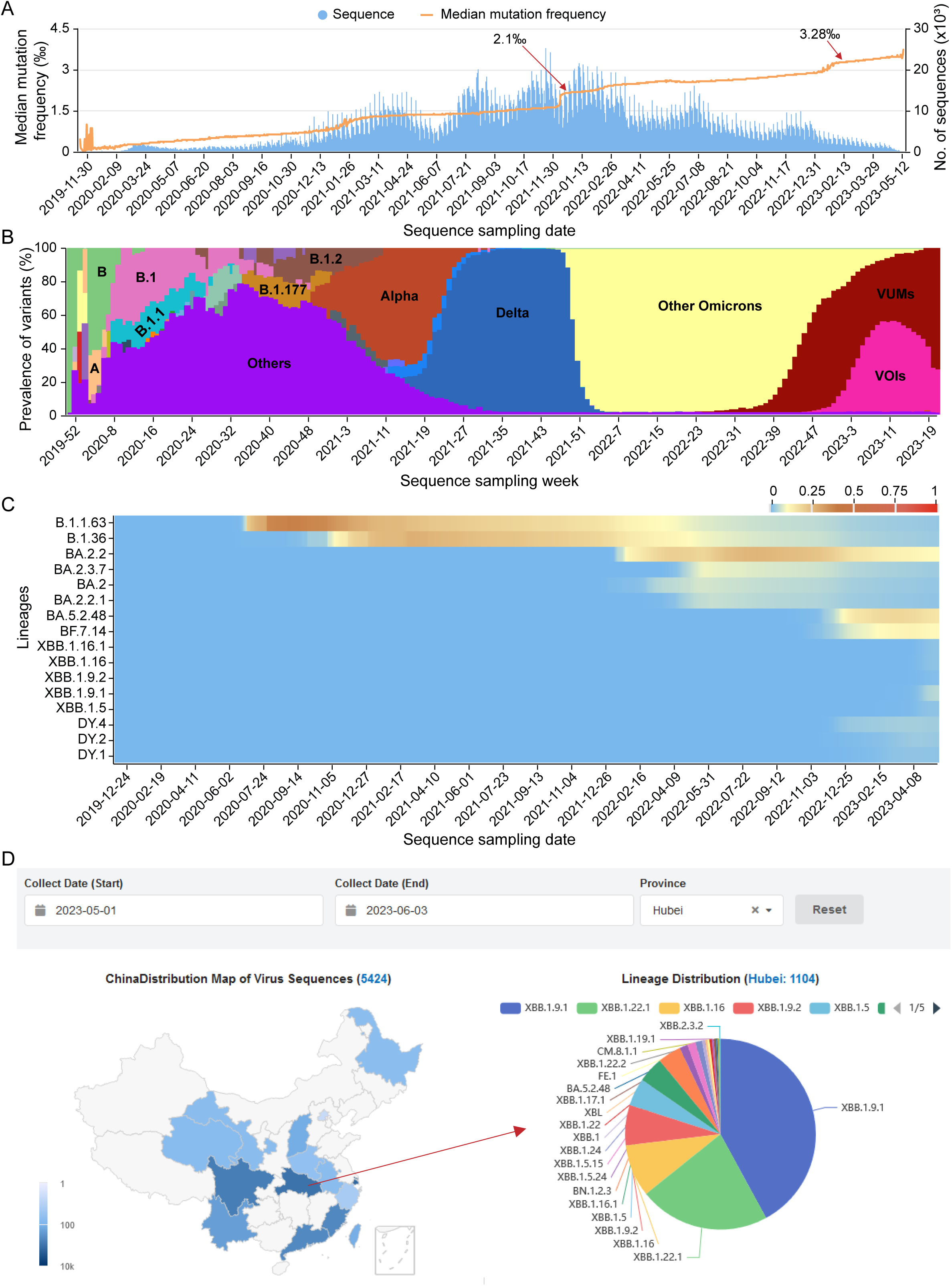
Monitoring of SARS-CoV-2 lineage evolution globally and regionally. **A**. Number of released sequences and the average mutation frequency along sequence sampling date. The mutation frequency is calculated by dividing the total mutation of each sequence by the genome length. **B**. The stacking diagram shows the proportion of top three prevalent Pango lineages or WHO define abbrevaitions used variants per week. **C**. Heatmap of the frequency of the cumulative sequences for selected lineage in China. **D**. Geographical distribution of the sequences number in China, and the pia chart shows the lineage proportions in provinces from May 1st to June 3 in 2023.

### Pre-warning of potential high-risk haplotypes and lineages

Early and accurate detection of potential high-risk SARS-CoV-2 haplotypes or lineages is a shared challenge for the scientific community in combating the virus. Leveraging the vast amount of genome sequences, we have developed a machine learning model called HiRiskPredictor[13] to predict potential high-risk haplotypes and update these predictions weekly in RCoV19. For each haplotype, a risk score ranging from 0 to 1 is calculated based on the available sequences at that time. Haplotypes with higher risk scores (> 0.5) are identified as potential high-risk haplotypes. A tabular table (**Figure 5A**) organizes the risk score, associated lineage, and transmission-related values (e.g., geographic entropy, betweenness, etc.) for each high-risk haplotype. Users can quickly search for specific haplotypes or lineages using different keywords, or sort the table by ‘Risk score’ to identify haplotypes with the highest risk scores. Additionally, a boxplot displays the top lineages (20 at most), ranked in descending order based on the median risk scores of all associated haplotypes. **Figure 5B** illustrates the prediction of 12 potential high-risk lineages as of May 31, 2023, with BN.1.2.3, XBB.1.5.24, XBB.1.9.1, XBB.1.16.1, and XBB.1.9.2 identified as the top five lineages. Importantly, the weekly predicted risk scores for all lineages are recorded, allowing users to track historical predictions to detect new warning lineages and understand their development trends (**Figure 5C**). Furthermore, the lineage prevalence, represented by the sequence proportion, is plotted to visualize global changes in epidemic variants (**Figure 5D**). For example, the dominant lineage XBB.1.5 accounted for 20% of all Omicron lineages but is gradually diminishing and being replaced by XBB.1.9.1.

**Figure 5.**
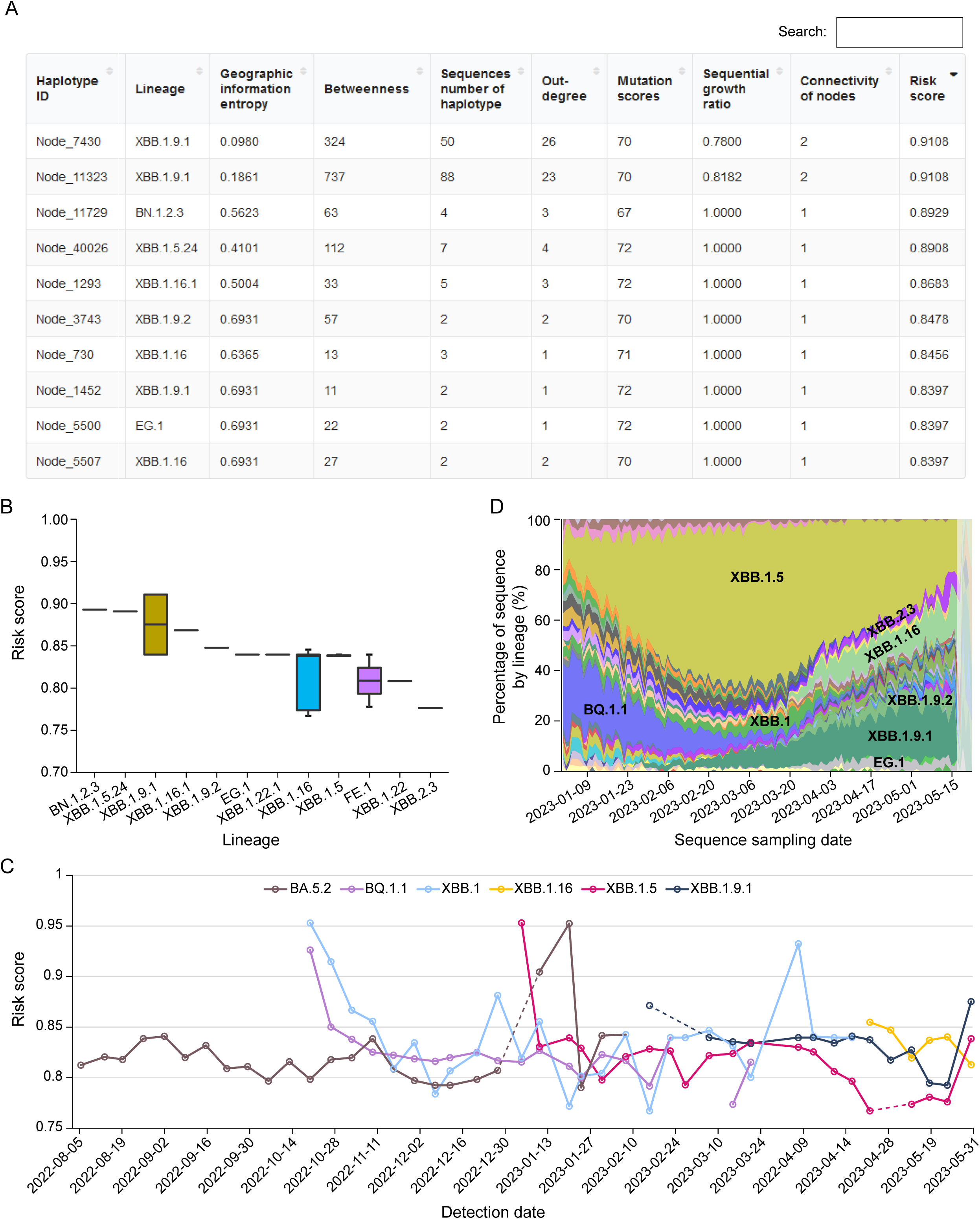
Pre-warning of potential high-risk haplotypes and lineages. **A**. A screenshot of the tabular table for all haplotypes with values of haplotype network features and its risk score. **B**. Boxplot of predicated risk score for all haplotypes of the top twenty lineages. As of May 31, 2023, 12 lineages have been predicted as potential high-risk lineages. **C**. Distribution of the historical risk scores for user selected lineages. **D**. Genomic prevalence of lineages based on sequence collection date.

These tools and visualizations provided by RCoV19 empower users to identify potential high-risk haplotypes and track the prevalence and evolution of lineages, contributing to early warning systems and informed decision-making in the fight against SARS-CoV-2.

### Mutation spectrum comparison between selected lineages or sequences

To facilitate the analysis of mutation spectra and comparisons between different lineages and sequences of SARS-CoV-2, we have developed two interactive modules within RCoV19. These modules allow users to explore mutation distributions and construct mutation maps on lineage level or sequence level.

In the inter-lineage or variants comparison module, users can examine the mutation patterns across WHO defined variants (e.g. Delta and Omicron) or Pango lineages (e.g. B.1.177 and XBB.1.5) and analyze mutations by genes or mutation frequency. For example, considering the top three prevalent lineages in the tenth week of 2023 (XBB.1, XBB.1.5, BQ.1) and previous VOCs (Alpha, Beta, Gamma, Delta, Omicron), it is evident that these lineages exhibit more mutations in the *S* gene (**Figure 6A**). Moreover, several novel mutations with high frequencies, such as S371F, T376A, and S477N (frequency > 0.89), have emerged in XBB.1 and XBB.1.5. Additionally, well-known mutations like D614G, known to enhance SARS-CoV-2 infectivity in human lung cells, and N501Y, associated with reduced vaccine protection in Delta, may explain the prevalence of ongoing XBB variants [19]. In addition to the extensively studied S protein, N protein mutations like R203K and G204R, implicated in increased transmission [20], are commonly observed in the top three ongoing lineages (**Figure 6B**). Notably, the N protein mutation P13L, which occurs at a high frequency of 90% in the top three ongoing variants, can significantly impair the CD8 T cell epitope (QRNAPRITF), leading to a loss of T cell recognition [21–23]. Similarly, amino acid deletions from position 31 to 33 in the N protein, with a high frequency of 90% among ongoing lineages, may contribute to improved replication efficacy or breakthrough infections, warranting further investigation in the future.

**Figure 6.**
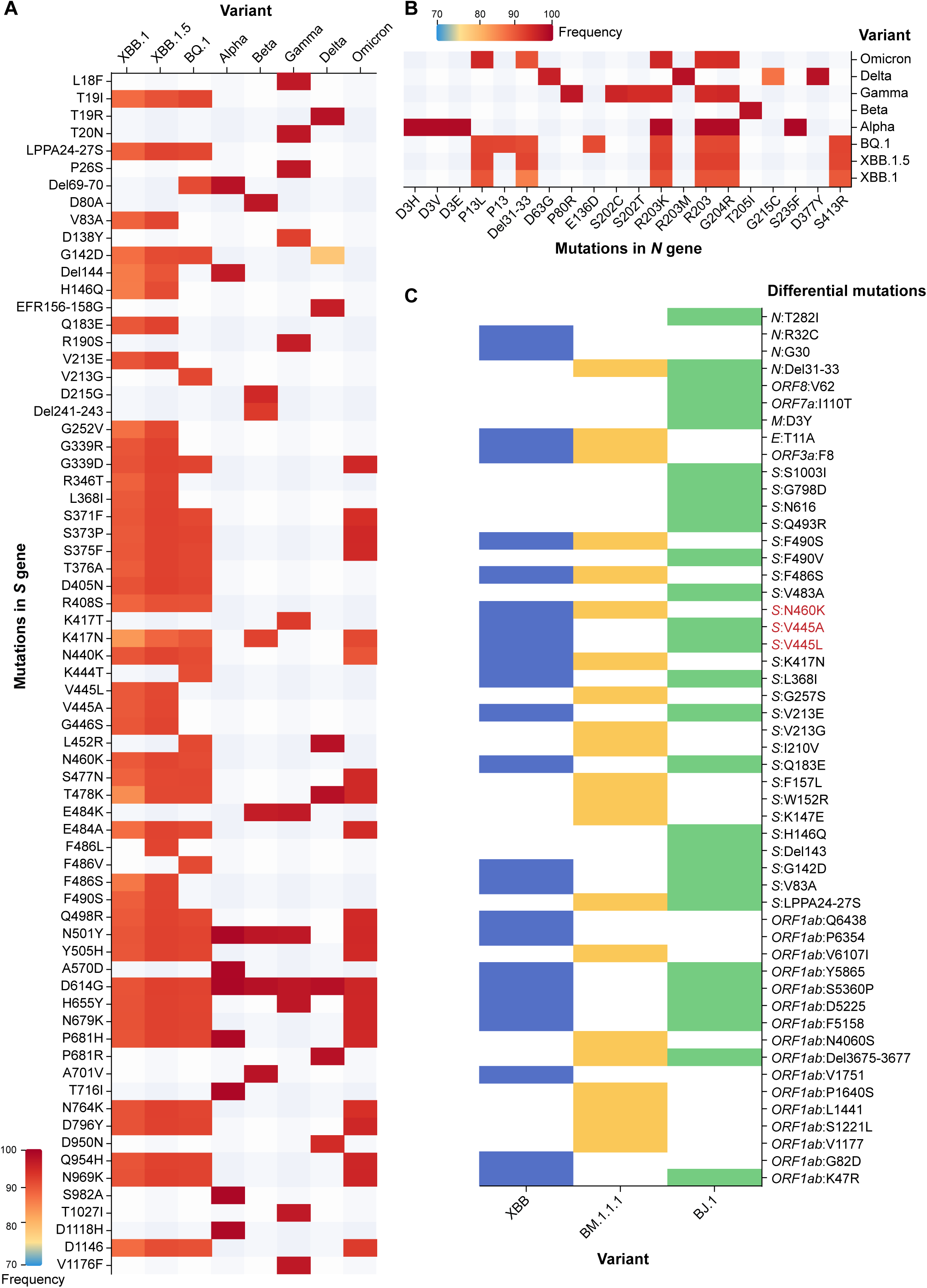
Mutation spectrum comparison among selected lineages and sequences. **A**. Lineage mutation comparison on *S* gene among top 3 prevalence lineages in 10^th^ week of 2023 (XBB.1, XBB.1.5, BQ.1) and previous VOC defined by WHO (Alpha, Beta, Gamma, Delta, Omicron) with mutation frequency. **B**. Lineage mutation comparison on *N* gene among top 3 prevalence lineages in 10^th^ week of 2023 (XBB.1, XBB.1.5, BQ.1) and previous VOC defined by WHO (Alpha, Beta, Gamma, Delta, Omicron) with mutation frequency. **C**. Sequence mutation comparison among sequences (XBB: EPI_ISL_15854782, BJ.1: EPI_ISL_14891585; BM.1.1.1: EPI_ISL_14733830) presented by differential mutations (refers to those after removing common mutations among sequences) in each sequence, the range between mutations in red color indicating possible recombination breakpoint.

In the multiple sequence comparison module, users can sensitively detect potential new lineages by comparing newly released sequences with the representative sequences of the latest lineages in our database. By inputting accession IDs and selecting the lineages of interest, the module displays a mutation matrix for comparison, which can be further refined interactively by genes or differential mutation sites. Additionally, the mutation matrix can be color-coded based on lineage, sampling date, or location. This module is particularly useful in narrowing down the breakpoint range in recombinant variants without the need for intensive sequence similarity calculations. For example, when analyzing the XBB recombinant lineage, comparing it with its parental sequences (BJ.1: EPI_ISL_14891585; BM.1.1.1: EPI_ISL_14733830) reveals that the breakpoint likely lies between V445 and N460 in *S* gene since XBB harbors V445 from BJ.1 and N460 from BM.1.1.1 (**Figure 6C**). Overall, this module complements existing platforms [24, 25] and aids in assessing the validity of newly assigned lineages.

These interactive modules within RCoV19 empower users to explore and compare mutation spectra across different lineages and variants, providing valuable insights into the evolution and characteristics of SARS-CoV-2 lineages.

### Investigation of the mutation effects on transmissibility and immune escape

A number of mutations have been confirmed to affect viral characteristics, including pathogenicity, infectivity, transmissibility, and antigenicity [8–10, 26, 27]. However, these knowledges are scattered across publications and always focuses on one aspect of a mutation or a variant. To facilitate the effective retrieval of mutation function, we have constructed an integrated knowledgebase by curating information from literatures and databases. Specifically, mutation knowledges are recorded and organized according to their impacts on infectivity/transmissibility, and effectiveness to antibodies, drug, and T cell epitopes.

Mutation-related information is collected and categorized based on their specific impacts. For each mutation, we have gathered details on its effects, including a comprehensive description, experimental methods used for characterization, and corresponding PubMed IDs (PMIDs) for reference. In the case of T cell epitope mutations, information on epitopes, HLA restriction, and corresponding T cell types has also been integrated. Overall, we have collected and summarized a total of 2696 single mutations, as well as other mutations such as SNPs and Indels, along with 19 combined mutations. Among these mutations, 76 affect infectivity/transmissibility, 131 are associated with drug resistance, 734 are related to antibody resistance, and 1817 mutations are located in T cell epitopes (**Figure 7A**). When considering the distribution of mutations across genes and open reading frames (ORFs), there is an uneven distribution. Specifically, in the S protein, 73 mutations (4%) have been reported to affect infectivity/transmissibility, while 733 mutations (58%) are associated with antibody resistance. This is understandable as the receptor-binding domain (RBD) of the S protein is responsible for virus binding to the ACE2 receptor and is a target for neutralizing antibodies. In the *ORF1ab* gene, 127 mutations (49%) are related to drug resistance, which may be attributed to *ORF1ab* being the target of most small molecule inhibitors.

**Figure 7.**
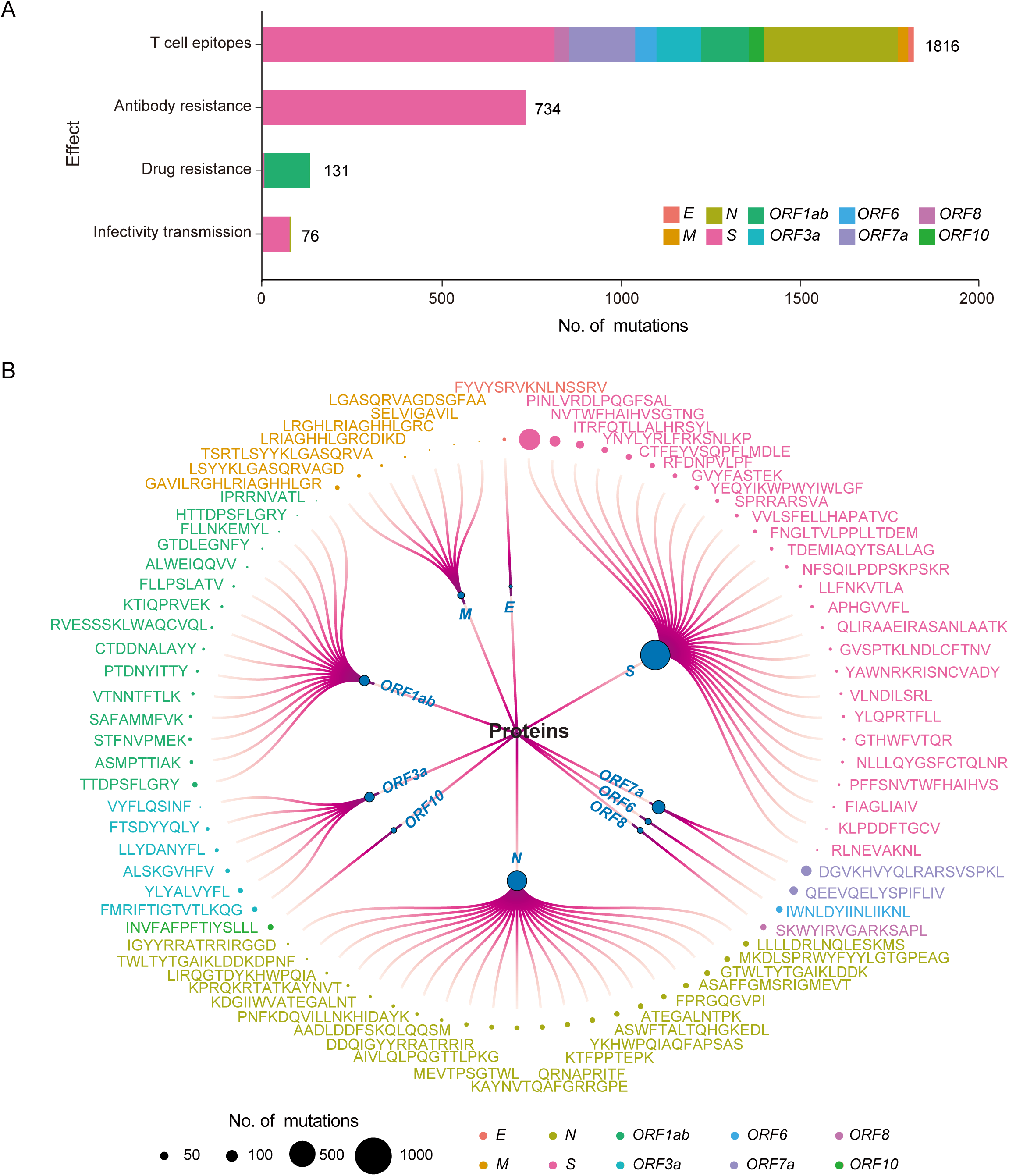
Mutation effects on SARS-CoV-2 viral characteristics. **A**. Collection of mutation effect knowledge. The horizontal axis represents the number of mutation types. **B**. Mutations occurring on experimentally verified T cell epitopes. The magnitude of the circles represents the number of mutations on each epitope, and different colors indicate T cell epitopes on different proteins.

Mutations located in T cell epitopes are of particular concern as they are dispersed across different proteins, posing challenges for the immune system to recognize and mount an effective response against various variants. Moreover, mutations in CD4 and CD8 T cell epitopes have the potential to disrupt HLA-peptide binding, leading to immune escape. The diverse epitopes found in different proteins exhibit distinct mutation patterns, which need to be carefully considered during the design of epitope-based vaccines (**Figure 7B**).

By providing a comprehensive and organized knowledgebase, researchers and users can easily access and retrieve information regarding the functional impacts of specific mutations. This integrated resource (https://ngdc.cncb.ac.cn/ncov/knowledge/mutation) enhances our understanding of the effects of mutations on viral characteristics and assists in the development of effective countermeasures against SARS-CoV-2 variants.

## Discussion

RCoV19 has been continuously updated and developed to support precise prevention of COVID-19. As an integrated repository for SARS-CoV-2 genome data, we have addressed various challenges by implementing a one-stop curation pipeline. This pipeline resolves issues such as sequence redundancy across different repositories, cross-linking between resources, and sequence quality evaluation. However, due to the lack of comprehensive clinical phenotype data, conducting in-depth association studies between massive genomic data and clinical outcomes, as well as unraveling the clinical significance of mutations, remains challenging. To enhance our understanding of disease spread and pathogenesis, we urge the collection and integration of clinical phenotype data of infected individuals to create a more comprehensive platform.

Timely monitoring and precise pre-warning based on genomic data are crucial for epidemic prevention. While there are platforms [4–6] available for spatiotemporal surveillance of mutations and variant evolution, there is a deficiency in platforms specifically focused on pre-warning of high-risk variants. In recent years, various machine learning-based prediction models have been proposed, such as PyR0 and VarEPS. PyR0, a hierarchical Bayesian multinomial logistic regression model, can identify mutations that are likely to increase SARS-CoV-2 fitness [28], while VarEPS evaluates the risk level of mutations and variants based on their transmissibility and affinity to neutralizing antibodies using a random forest model [7]. In RCoV19, we have developed a LightGBM model called HiRiskPredictor [13], which calculates a comprehensive risk score and predicts potential high-risk haplotypes on a weekly basis. In the future, we aim to provide multidimensional pre-warning by combining the strengths of different AI models and features.

Genetic mutation spectra play a critical role in determining the virological characteristics of different virus strains. Sequence comparison remains the primary approach for identifying differences in mutation spectra. Although the Pango dynamic phylogeny-informed nomenclature system has made significant contributions to tracking genetic diversity and classifying SARS-CoV-2 lineages, there is often a time gap before sporadic variants occurring in specific regions are designated as new lineages. To stay updated on SARS-CoV-2 mutations more sensitively and identify novel lineages earlier, RCoV19 now supports the comparison of newly released sequences with representative sequences of the latest lineages. This feature complements existing public platforms [24, 25] and assists in verifying the assigned lineages of newly released sequences.

Numerous mutations have been identified that can increase the severity of infections, enhance transmissibility, and enable evasion of natural and vaccine-induced immunity [29]. Through comprehensive literature curation, we have consolidated a wealth of knowledge regarding the effects of mutations on viral infectivity, resistance to antibodies and therapeutic drugs, and alterations to T cell epitopes. However, further investigation is needed on mutations that impact disease severity. For example, the mutation S194L in the nucleocapsid (N) protein has a notably high frequency among individuals with severe clinical manifestations [30], suggesting its potential contribution to disease progression. Additionally, most of the knowledge on mutation effects is curated from published literature or databases. Future improvements could focus on structural bioinformatics-based prediction of mutation effects, which would enhance our understanding of future pandemics and aid in the development of preventive measures and treatment strategies. In conclusion, knowledge of mutation effects is essential for effective public health interventions, the development of therapeutics, and the creation of pre-warning models.

## Methods

### Pre-warning of potential high-risk haplotypes

All the complete and high-quality SARS-CoV-2 sequences and metadata in RCoV19 were used to predict potential high-risk haplotypes weekly. First, we calculated the population mutation frequency (PMF) for each mutated site within every month. Then, those non-UTR mutations with PMF > 0.005 were selected for haplotype network construction by McAN with default parameters. Next, the result of the haplotype network was loaded into HiRiskPredictor with a pre-trained machine learning algorithm (LightGBM) to perform the forewarning analysis process. The HiRiskPredictor automatically extracts features, such as out degree, geographic information entropy, betweenness, etc., for each haplotype in the network. And HiRiskPredictor infers a risk score indicating the likelihood of a haplotype being positive or classified as high-risk according to those features via the pretrained model. If the predicted risk score of a haplotype is greater than 0.5, it is defined as a high-risk haplotype.

### Mutation spectrum comparison between selected lineages or sequences

Only complete and high-quality genome sequences that have been previously evaluated were employed for the following sequence comparison. To achieve this, the genome sequences were aligned using MUSCLE (version 3.8.31) [15034147] and compared against the initial SARS-CoV-2 genome release (GenBank: MN908947.3). The identification of sequence variations was accomplished using a custom Perl program. At lineage level, mutations among all complete and high-quality sequences of selected variants are displayed in heatmap with customized population mutation frequency. At sequence level, newly emerged sequences with fixed mutations in specific lineage are chosen as representative sequences. After compared with reference genome, mutations among different sequences are displayed in heatmap. Instead of representative sequences, this module also supports to conduct sequence comparison according to Input Sequence Accession within the database.

### Investigation of the mutation effects on transmissibility and immune escape

Through a comprehensive literature curation, we have collected a curated list of epitopes that have been experimentally validated. These experiments involved interferon-γ (IFN-γ) enzyme-linked immunospot (ELISpot) assays, complex class I (pMHCI) tetramer staining, and peptide-stimulated activation-induced marker (AIM) assays .etc. Subsequently, we employed an in-house program to integrate all available mutation data across the genome with those effective epitopes and filter mutations with sequences account lower than 2000. Following this, we have conduct a more precise literature curation to search for mutation effect occurring on epitopes to illustrate their functions in T cell recognitions.

## Data availability

SARS-CoV-2 genomes, mutations in vcf/tab format and their annotations are publicly available at https://ngdc.cncb.ac.cn/ncov/.

## CRediT author statement

**Cuiping Li**: Methodology, Formal analysis, and Writing - Original Draft. **Lina Ma**: Data curation, Methodology, and Writing - Original Draft. **Dong Zou:** Software. **Rongqin Zhang**: Data curation, Methodology, and Writing - Original Draft. **Xue Bai**: Data curation, Writing. **Lun Li**: Methodology. **Gangao Wu**, **Tianhao Wu**, **Wei Zhao** and **Enhui Jin**: Data curation. **Yiming Bao**: Conceptualization, Supervision, and Writing - Review & editing. **Shuhui Song**: Conceptualization, Methodology, and Writing - Review & editing. All authors have read and approved the final manuscript.

## Acknowledgments

This work was supported by grants from the National Key R&D Program of China (Grant No. 2023YFC3041500, 2021YFF0703703), the Key Collaborative Research Program of the Alliance of International Science Organizations (ANSO-CR-KP-2022-09), National Natural Science Foundation of China (Grant No. 32270718), Beijing Nova Program (Z211100002121006) and the Youth Innovation Promotion Association of Chinese Academy of Sciences (Grant No. Y2021038, 2019104). Thanks to all colleagues who participated in data curation and provided valuable suggestions. We also thank a number of users and CNCB members for reporting bugs and sending comments. Complete genome sequences used for analyses were obtained from the Genome Warehouse (GWH) and GenBase of CNCB-NGDC, CNGBdb, GenBank, GISAID, and NMDC resources. The construction of the knowledge information refers to the COG-UK-ME, CoV-RDB, SCoV2-MD, and ESC databases. We acknowledge all sample providers and data submitters.

