## Supplementary figures and images for "RCoV19: A One-stop Hub for SARS-CoV-2 Genome Data Integration, Variants Monitoring, and Risk Pre-warning"

### Supplemental figure 1

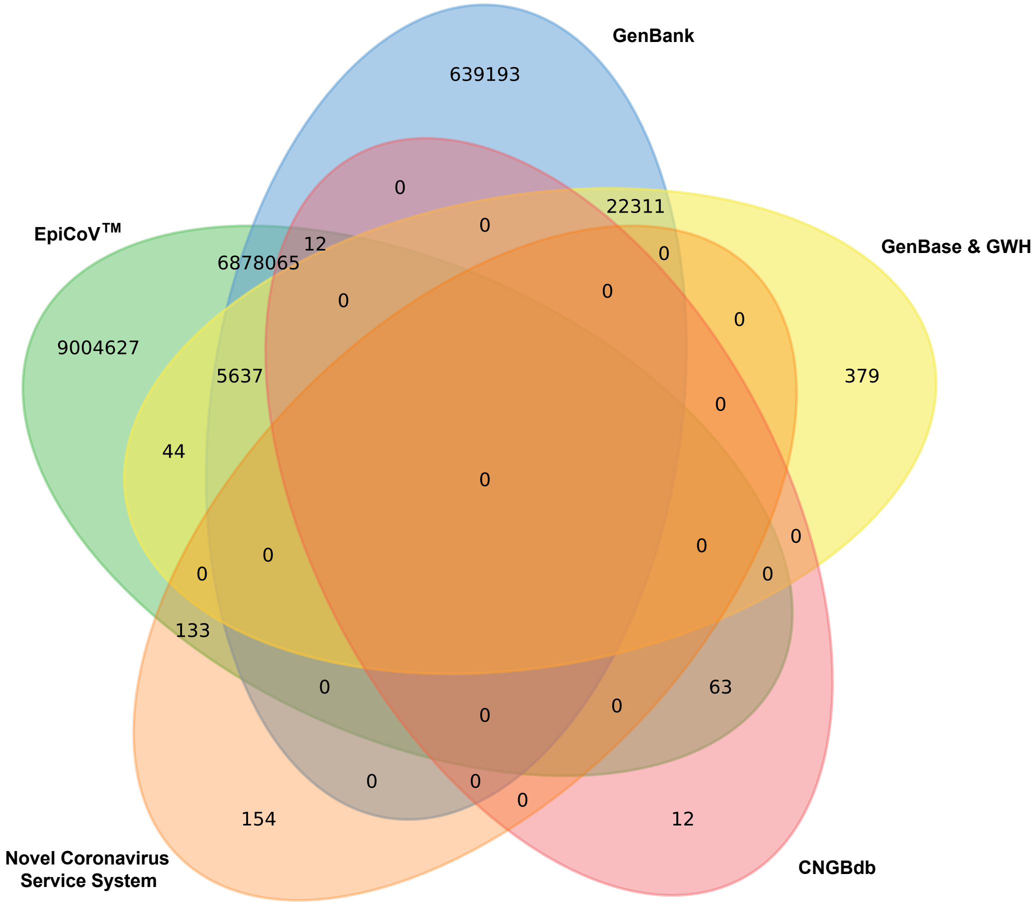
